# Comparative Analysis of the Microbiota of *Vaccinium myrtillus* and *Vaccinium uliginosum* in the Central Italian Apennines

**DOI:** 10.1101/2025.05.26.655108

**Authors:** Francesca Vaccaro, Ginevra Cambi, Stefano Biricolti, Edgardo Giordani, Alessio Mengoni, Camilla Fagorzi

## Abstract

*Vaccinium* is a genus of shrubs known for their antioxidant and anti-inflammatory compounds. While the commercially cultivated blueberry (*V. corymbosum*) originates from North America, European species like *myrtillus* (bilberry) and *V. uliginosum* (bog bilberry) remain uncultivated due to poor adaptability to conventional growing media. At present, bilberries, which are more valued for fresh consumption, are exclusively harvested from wild populations, particularly in subalpine and mountain environments, where unique soil and climate conditions prevail.

This study explores the bacterial and fungal communities associated with *V. myrtillus* and *V. uliginosum* in wild populations of the Central Italian Apennines, using 16S rRNA and ITS amplicon sequencing. Microbiota from plant and soil were analyzed, revealing distinct microbial community compositions based on species and plant compartments. Bacterial diversity was highest in bulk soil, while fungal diversity dominated plant tissues. Co-occurrence network analysis showed greater connectivity in *V. uliginosum* microbiota, suggesting higher resilience. Functional predictions indicated roles in nitrogen cycling, cellulose degradation, and plant-microbe interactions. These findings offer insights into the native microbiota of wild *Vaccinium* species and could inform conservation and cultivation efforts.

## 1. Introduction

*Vaccinium spp*. sect. Cyanococcus are perennial flowering plants of the Ericaceae family, known for their blue or purple berries. They encompass approximately ∼500 shrubs/small tree species globally ^1^. Commercial blueberry cultivars are all native to North America and mostly derive from breeding programs of blueberry (*V. corymbosum* L.) and cranberry (*V. macrocarpon* Ait.) ^2^. Blueberry fruits have high content of flavonoids, phenolic acids, polyphenols, anthocyanins, pro-anthocyanidins, and stilbenes, and show antioxidant and anti-inflammatory activities (see for instance ^3^). In the last years, numerous studies have investigated the role of blueberry fruit juice on human health and human gut microbiota ^4,5^ (REF). In the European flora *Vaccinium* species are represented by four species only, *V. myrtillus, V. uliginosum, V. oxycoccus* and *V. vitis idaea* ^6^. The bilberry or forest blueberry *V. myrtillus*, and the lingonberry, *V. vitis idaea*, are the only two of any economic significance, but are mainly consumed locally, within traditional and family-based practices. Forest bilberry (*V. myrtillus*, L.) is a small deciduous, broadleaf shrub with oval, light green leaves; the flowers are white with fused petals. In Italy, it is found in the Alps and the Apennines, particularly in subalpine and montane beech forests (from 500 to 2500 meters). The genetic diversity of the Tuscan mountain populations of bilberry and the biochemical traits of both *V. myrtillus* and *V. uliginosum* have been clearly shown ^7^. *Vaccinium* species naturally grow in well-drained, sandy acidic (pH 4.0-5.5) soils that tend to be low in nutrients and with 2-7 % of organic matter ^8^. Moreover, in contrast to cultivated blueberries, forest bilberry of the Tuscan mountains, notwithstanding the recent results achieved in the *in vitro* and *in vivo* propagation ^9^ shows a very low adaptability to natural soil and growing media specifically selected well suited for other Ericaceae species, including blueberry (*V. corymbosum*).

Although tolerant of partial shade, these plants require sufficient sunlight to maximize fruit production, but thrive in cool, humid climates and can be damaged by late frosts, drought, or excessive heat. In Central Italy, populations of forest bilberry are consequently particularly threatened as a result of climate change. Moreover, in Central Italian Apennines *V. myrtillus* and *V. uliginosum* often co-occur and in some cases *V. uliginosum* is invading the spots of *V. myrtillus*, creating additional concern over the preservation of *V. myrtillus* populations ^7^. Since the soils in which these plants grow are not typical, compared to the soils in which most crops are grown, the rhizosphere microbiota assumes somewhat unique features, especially in relation to the contribution to plant nutrition in low organic matter, acidic soils. Indeed, the bacterial communities in the rhizosphere of *Vaccinium* species are dominated by various groups, as Proteobacteria and Acidobacteria in North American cultivated *Vaccinium* species ^10^ or dominated by Actinobacteria (for the Asian species *V. ashei* ^11^ or by all these three bacterial groups ^12^. These bacteria are likely involved in various possible host beneficial processes such as nitrogen cycling, including nonsymbiotic nitrogen fixation, phosphorus solubilization, and production of plant growth-promoting hormones. Phosphate-solubilizing bacteria should be particularly important in the acidic soils where *Vaccinium* species are typically found, as phosphorus availability is often limited under such conditions. Fungi are another essential component of the rhizosphere microbiome in *Vaccinium* species. Mycorrhizal fungi, particularly ericoid mycorrhizae (ErM), are of special interest ^13^. ErM are fungi that form symbiotic relationships with the roots of *Vaccinium* plants, enhancing their ability to absorb water and nutrients, especially phosphorus. The symbiosis between *Vaccinium* species and ErM is highly specialized, with *Vaccinium* roots providing carbon to the fungi in exchange for improved nutrient uptake. Recent findings also suggest that the colonization by ErM fungi could also involve above-ground tissues, such as stem and leaves ^14^, possibly opening to speculate about more direct modulation of secondary metabolite production by the plant. The interaction between ErM and bacteria seem to have a role in resilience to heat stress ^15^. The co-occurrence and invasion of *V. uliginosum* of the spots of *V. myrtillus* can lead to hypothesis that differences in the recruitment of the associated microbiota from the soil could be relevant for thriving under the pedo-climatic conditions of these areas. Indeed, comparative studies on *V. uliginosum* and *V. myrtillus* microbiota are lacking, while only a few studies analyzed some aspects of the associated microbiota of these two species individually ^14,16,17^. Moreover, the still lack of cultivation practices for *V. myrtillus* could be partly attributed to challenges in establishing a suitable microbiota under cultivation. It is then interesting, both for understanding the presence of species-specific features of the forest and bog bilberry microbiomes and defining specificities for approaching bilberry cultivation, the analysis of the rhizosphere and endosphere microbiota of *V. uliginosum* and *V. myrtillus* collected in the wild spots, either in single or co-occurring populations.

Here we report findings obtained from microbiota analysis performed by amplicon sequencing of 16S rRNA gene (bacteria) and Internal Transcribed Spacer (ITS, fungi) for five spontaneous populations in Central Italian Apennines. We aim to identify the importance of site and host species (*V. uliginosum* and *V. myrtillus*) in affecting the diversity and taxonomy of bacterial and fungal microbiota, trying to hypothesize also possible roles of the microbiota in the partial invasiveness by *V. uliginosum* of the spots colonized by *V. myrtillus*.

## 2. Results and Discussion

### 2.1 Overall Diversity of the Bacterial and Fungal Microbiota

For the V3-V4 amplicon sequencing, a total of 4’159’967 16SrRNA amplicons reads were obtained, and 3’836’541 (92% of total reads) passed quality filtering. After the merging step and chimera removal, a total of 2’194’373 reads were obtained (Supplementary Information File, Table S1), corresponding to 1921 ASVs (Supplementary Dataset). For the ITS amplicon sequencing a total of 4’138’283 ITS reads were obtained, and 3’727’259 (90% of total reads) passed quality filtering (Supplementary Information File, Table S2). After the merging step and chimera removal, a total of 3’537’840 reads were obtained, corresponding to 728 ASVs (Supplementary Dataset). Rarefaction curves obtained from the ASVs reached a plateau for all samples (Supplementary Information File, Figure S1), indicating a satisfactory survey of the bacterial diversity (Good’s coverage, Supplementary Information File, Tables S3 and S4), which allowed to estimate alpha diversity indices.

### 3.2. Bacterial and fungal microbiota are differentially affected by host plant species

Alpha diversity estimates were obtained (Figure 1), with comparisons made between different plant compartments (origin), plant species, and sampling sites for both bacterial (16SrRNA) and fungal (ITS) microbiota. The bacterial microbiota diversity analysis in *V. myrtillus* and *V. uliginosum* across compartments (leaves, roots, rhizosphere) revealed significant differences in plant-associated microbiota compared to bulk soil. In particular, Shannon and Chao1 indices showed the highest values for bulk soil, followed by rhizosphere and roots, and then by leaves. Interestingly, an inverse trend was observed for Pielou’s evenness values, leaves having the highest, followed by root, and then soil and rhizosphere. The analysis across species did not show differences between the whole *V. myrtillus* and *V. uliginosum* samples but still showed bulk soil having different alpha diversity values compared to plant-associated samples. For the fungal microbiota, alpha diversity analysis (Figure 1) revealed, in contrast with bacterial microbiota, higher Pielou’s evenness in leaves compared to the other compartments. Additionally, while bacterial microbiota alpha diversity was similar between V. *myrtillus* and *V. uliginosum*, the fungal microbiota alpha diversity was higher in *V. uliginosum* samples compared to *V. myrtillus* for Shannon and Simpson indices.

**Figure 1.**
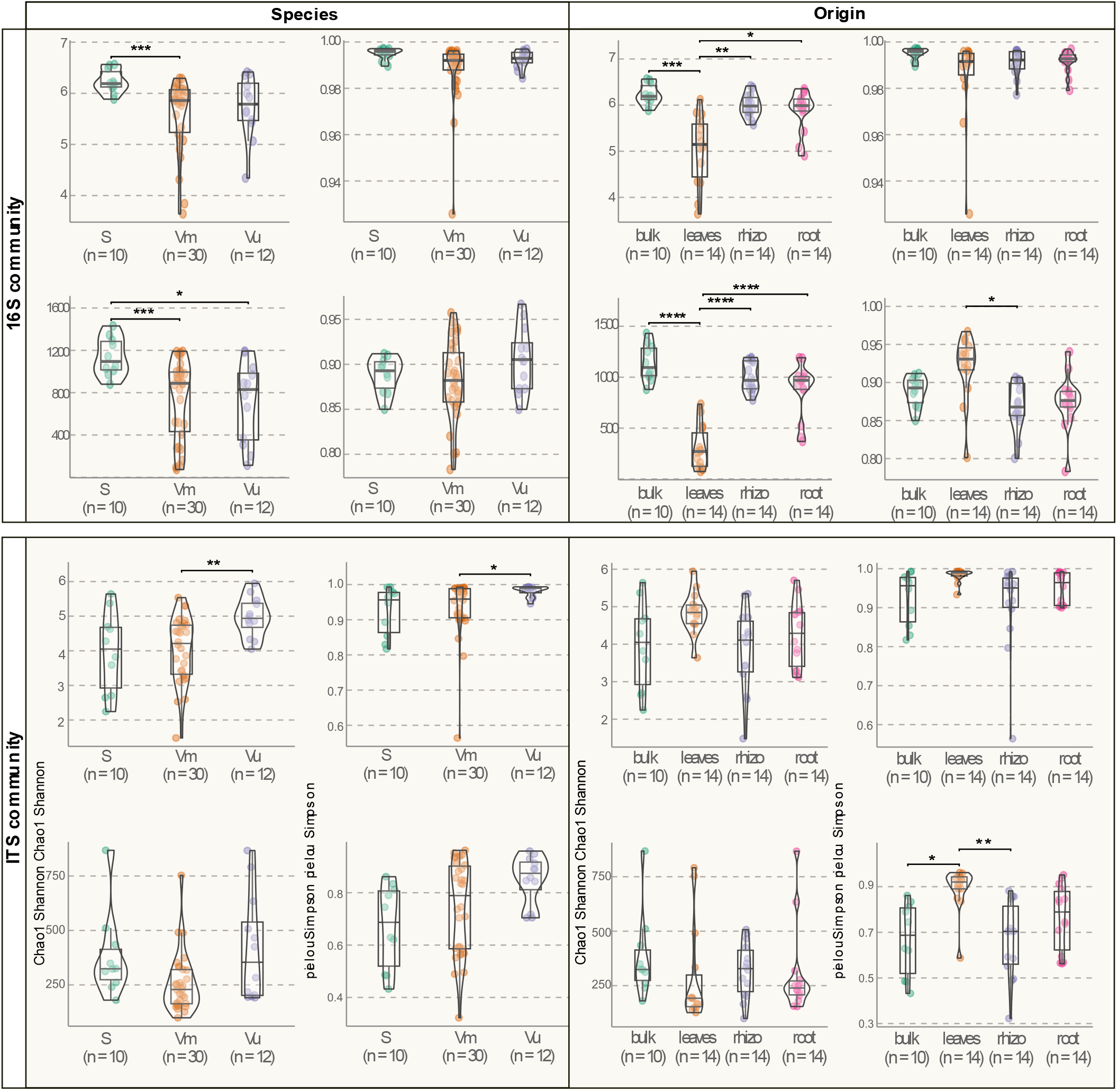
Alpha diversity indices of the bacterial and fungal microbiota associated with *V. myrtillus* and *V. uliginosum* (a) Shannon index; (b) Simpson index; (c) Chao1 index; (d) Pielou index). Lines and the numbers above the lines indicate the p-values of the contrasts (Wilcoxon test). Bulk soil (S), *V. myrtillus* (Vm), *V. uliginosum* (Vu). Asterisks indicate level of significance (* p≤ 0.05; **p≤0.01; *** p≤ 0.001; **** p≤ 0.0001).

The evidence reported above on alpha diversity of bacterial and fungal microbiota suggests that these two fractions of microbiota are differentially affected by the host plant. In fact, while for the bacterial fraction the host plant clearly selects a group of taxa (Shannon and Chao1 diversity is lower than that of soil), the fungal fraction showed that leaves harbor higher diversity than soil. Moreover, the fungal fraction also showed between host species differences. Such different behaviors of bacterial and fungal plant-associated microbiota with environmental changes (including site, host plant genotypes etc.) have been observed frequently in other systems. In *Photinia* x *fraseri* we previously observed that the fungal leaf microbiota is more influenced by plant growth conditions (presence of particulate pollution) than the bacterial microbiota ^18^. Similar results have been reported for several plant species in relation to drought stress ^19^. Indeed, bacterial and fungal communities associated with plants exhibit distinct sensitivity to environmental conditions, influenced by factors such as water availability, land use, and soil properties ^20–22^. Our results are adding novelty to this evidence; overall, the body of results on alpha diversity indicates that both V. *myrtillus* and *V. uliginosum* actively shape their microbiota, possibly resulting in communities that are potentially functionally specialized, either among plant compartments or between host species. The indication that the fungal fraction is able to differentiate between the two species is stirring attention over the possibility that species-specific associations between fungi and host plant could be key in shaping the microbiota.

A similar pattern of differences was observed for the taxonomic similarities among microbiota (Figure 2). NMDS showed a clear separation between leaf and bulk/rhizosphere associated bacterial fraction, with samples from root compartment being more widely dispersed between these two poles, suggesting greater taxonomic heterogeneity in the root compartment microbiota. PERMANOVA (Table 1) confirmed that both host species and origin (compartments leaves, root, rhizosphere, and bulk soil) significantly influenced bacterial composition (p < 0.05), with a significant interaction effect (p = 0.003). For the fungal community (Figure 2b) the NMDS did not show clear clusterization, possibly suggesting that the fungal taxa identified could be relatively ubiquitous in different plant compartments and between *V. myrtillus* and *V. uliginosum*. However, the PERMANOVA results indicated that host plant species and origins (plant compartments) do have a statistically significant effect on fungal microbiota composition, in line with also the results from alpha diversity. The relationship between fungal taxonomic diversity and host plants was further confirmed by a beta-dispersion test (Table S1).

**Table 1.** Microbial communities are structured by *Vaccinum* species, sampling site, and compartment. Permutational Multivariate Analysis of Variance (PERMANOVA performed to assess the effect of origin and site on beta diversity based on Bray-Curtis distances. The model tested the interaction between Species, Site, and Origin (Species x Site x Origin) for both (a) the bacterial (16SrRNA) and (b) fungal (ITS) microbiota. Asterisks indicate significance thresholds (*, <0.05; ** <0.01; *** < 0.001).

|  | a) Bacterial microbiota |  | b) Fungal microbiota |  |
| --- | --- | --- | --- | --- |
|  | F | Pr(>F) | F | Pr(>F) |
| Origin | 1.5917 | 0.010 ** | 1.8337 | 0.001 *** |
| Species | 1.6627 | 0.006 ** | 1.8197 | 0.002 ** |
| Site | 1.8056 | 0.013 * | 1.9725 | 0.002 ** |
| Origin x Species | 1.8699 | 0.002 ** | 1.2567 | 0.074 |
| Origin x Site | 1.9858 | 0.001 *** | 1.802 | 0.001 *** |
| Species x Site | 1.756 | 0.004 ** | 0.9642 | 0.514 |
| Origin x Species x Site | 1.6658 | 0.002 ** | 1.9498 | 0.001 *** |

**Figure 2.**
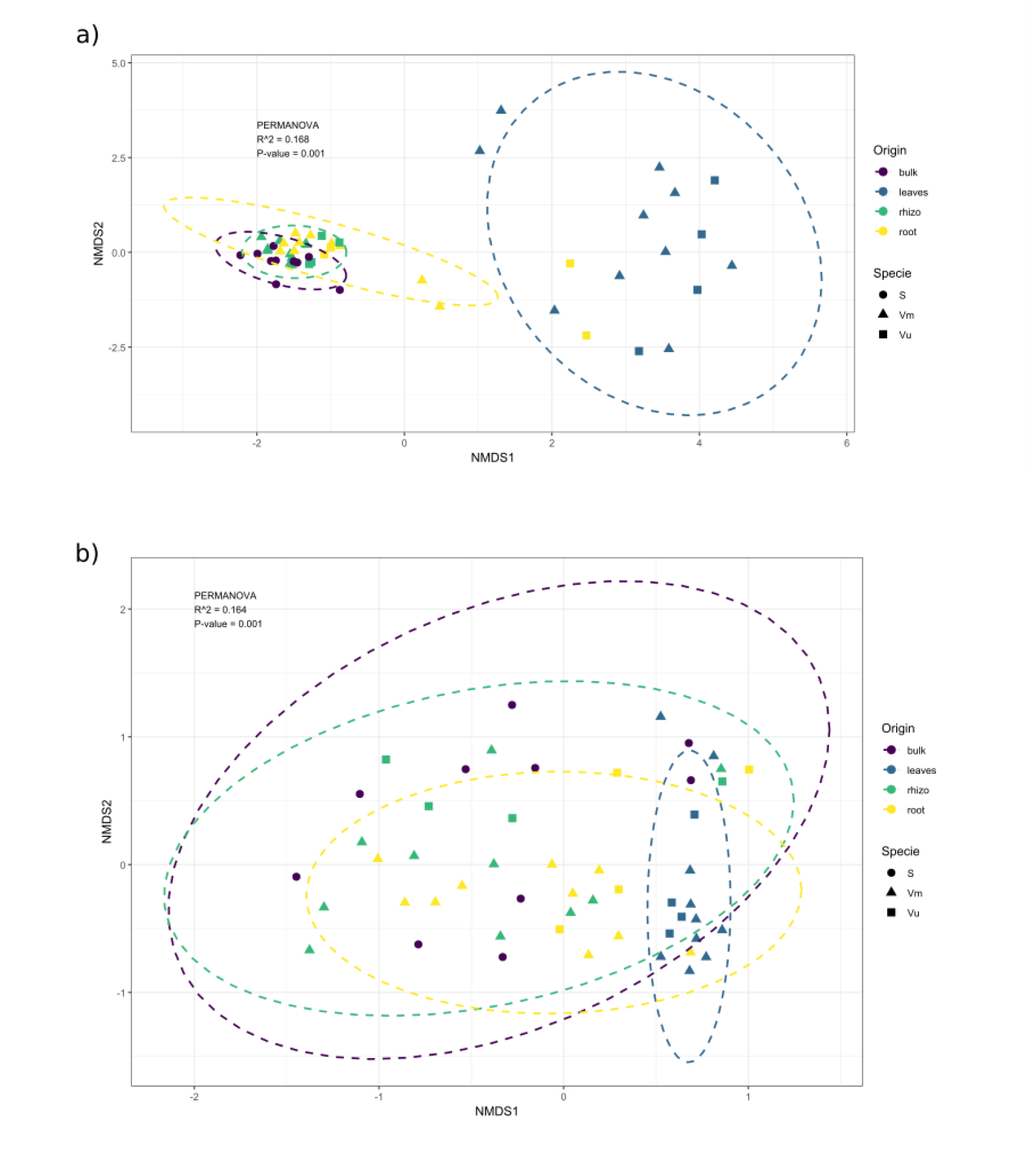
Nonmetric multidimensional scaling (NMDS) of a) bacterial and b) fungal community. Ordination based on Bray-Curtis distance. Colors indicate origin of the samples (violet= bulk soil, blue=leaves, green=rhizosphere, yellow=roots). Shape indicates species (triangle= *V. myrtillus*, square= *V. uliginosum*, circle= bulk soil.

Actually, it is known that the fungal microbiota associated with *Vaccinium* species varies significantly, influenced by several factors such as plant species (genotype), environmental conditions, and soil characteristics. For the rhizosphere microbiota, for instance, research comparing the rhizosphere microbiomes of southern highbush blueberry (*Vaccinium corymbosum*), Darrow’s blueberry (*Vaccinium darrowii*), and rabbiteye blueberry (*Vaccinium virgatum*) revealed significant differences in the abundance of ErM fungi among these species. ErM fungi are indeed important for the adaptation to acidic soils with low nutrient availability ^23,24^. For *V. myrtillus* and *V. uliginosum* previous studies have actually found relevant differences in the endophytic fungal microbiota (Daghino et al., 2022; Yang et al., 2018).

We cannot exclude that these differences in fungi can impact the ability of *V. myrtillus* and *V. uliginosum* to adapt to specific habitats and explain the partial invasion of *V. myrtillus* by populations of *V. uliginosum*. However, we cannot exclude that the statistically significant differences detected could be influenced by the disparity in within-group variability. Actually, unequal sample sizes, particularly for *V. uliginosum*, could lead to greater variability in dispersion, potentially distorting the interpretation of species-related differences.

### 2.3 Taxonomic characterization of V. myrtillus and V. uliginosum microbiota

The taxonomic representation of *V. myrtillus* and *V. uliginosum* showed among the most abundant bacterial classes Alphaproteobacteria, Bacteroidia, Acidobacteriae and Actinobacteria, and for fungi Agaricomycetes, Sordariomycetes, Dothideomycetes, Leotiomycetes, Tremellomycetes, Mortierellomycetes and Glomeromycetes were the most abundant classes (Figure S1). Leaves vs. bulk soil bacterial microbiota showed the highest statistically significant differences (by DESeq2) in the presence and abundance of ASVs affiliated with several classes (Figure 3a). Enrichment in Bacteoridia and Alphaproteobacteria was shared among all the tested contrasts. Interestingly *V. myrtillus* and *V. uliginosum* contrast showed statistically significant differences in 7 classes (more abundant in *V. myrtillus* than *V. uliginosum*). A previous study in *V. angustifolium* the rhizosphere was found to be predominantly inhabited by Alphaproteobacteria, mainly from the genus *Bradyrhizobium* ^24^. The role of these bacteria (nitrogen-fixer of leguminous plants) is still unclear but is in line with a general observation that Alphaproteobacteria are a key bacterial class in plant-microbe interaction ^26^. The same and another study in *Vaccinium corymbosum* ^27^ found also abundant presence of Acidobacteria in the rhizosphere suggests their potential role in soil nutrient cycling. It is worth noticing in our study that Bacteoridia differentiate among either species or origin. Although Bacteoridia are well studied as an important member of the human intestinal tract, their functional roles in plant microbiomes remain largely elusive ^28^. However, recent research has shown that they actively colonize various plant districts and can contribute to nutrient cycling ^28,29^. As for the bacterial microbiota, several fungal classes showed statistically significant enrichment between plant samples and soil and between *V. myrtillus* and *V. uliginosum* (Figure 3b) as: Agaricomycetes, Dothideomycetes, Leotiomycetes, Mortierellomycetes, Saccharomycetes, Sordariomycetes and Tremellomycetes. The presence of these fungal classes in the microbiota of *Vaccinium* spp. across different compartments (roots, rhizosphere, and leaves) could be justified based on their ecological roles and their interactions with plants in these specific niches. While **mycorrhizal fungi** (like **Leotiomycetes** and **Glomeromycetes**) enhance nutrient availability in the root zone by facilitating nutrient exchange between plant roots and the soil ^14,30^, **saprotrophic fungi** (such as **Tremellomycetes, Mortierellomycetes**, and **Agaricomycetes**) break down organic matter, releasing nutrients into the soil and enhancing plant health ^31^. In previous studies, while in *V. myrtillus* fungi from the genus *Hyaloscypha* and the *Phialocephala-Acephala applanate* complex, in *V. uliginosum* genera as *Rhizoscyphus* and *Meliniomyces* dominated ErM communities, followed by *Clavaria, Oidiodendron, Lachnum, Acephala*, and *Phialocephala* were recovered ^25,32^. All (but *Clavaria*) of these taxa are belonging to Leotiomycetes, one of the above reported classes, which includes many ericoid mycorrhizal fungi. Clavaria is a member of Agaricomycetes, another of the classes which statistically differentiates between *V. myrtillus* and *V. uliginosum*.

**Figure 3.**
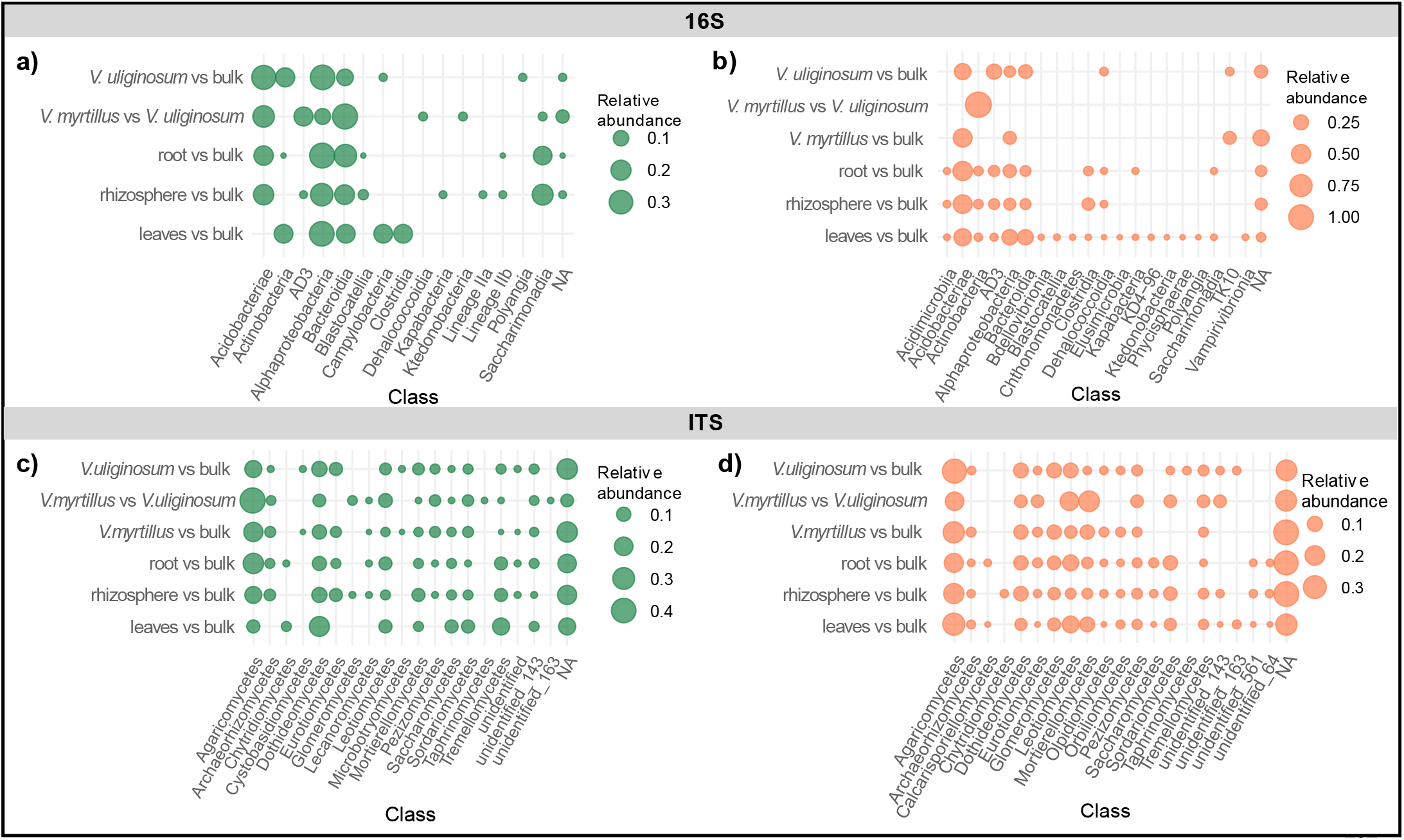
Taxonomic enrichment between microbiota. Bubble plot depicting the relative abundance of classes for a) the bacterial (16S) microbiota and b) the fungal (ITS) community, based on differentially abundant ASVs (padj<0.05) across each contrast. Diameter of bubbles is proportional to relative abundance of each class. “Bulk” indicates bulk soil samples.

### 3.4. Network analysis of interactions within the microbiota

To further inspect the microbiota of *V. myrtillus* and *V. uliginosum* in relation to host-species differences, co-occurrences among the total set of ASVs (both bacterial and fungal) were computed and network representations were used to determine the presence of microbial groups which may potentially interact among them. Such representation may help to decipher the structure of microbiota to help understanding the ecological rules guiding community assembly ^33^. Results from analysis of co-occurrence networks (Figure 4) showed three clearly different patterns among soil, *V. uliginosum* and *V. myrtillus*. As expected (Yang et al., 2024), soil microbiota showed the highest number of edges (7947) when compared to the plant-associated microbiota, indicating a very high level of interactions between microbial phyla. The high connectivity of soil co-occurrence network highlights the importance of native soil in providing many ecological niches for microorganisms, aligning with previous studies emphasizing the role of soil microbiota in maintaining ecosystem stability and functioning ^35^. The relatively lower number of nodes (284) compared to the number of edges suggests that the distinct microbial groups interact extensively with each other, forming a dense network and describing a possible high degree of ecological complexity within the soil. This result may suggest good resilience and ecosystem services provided by the samples soil microbiota.

**Figure 4.**
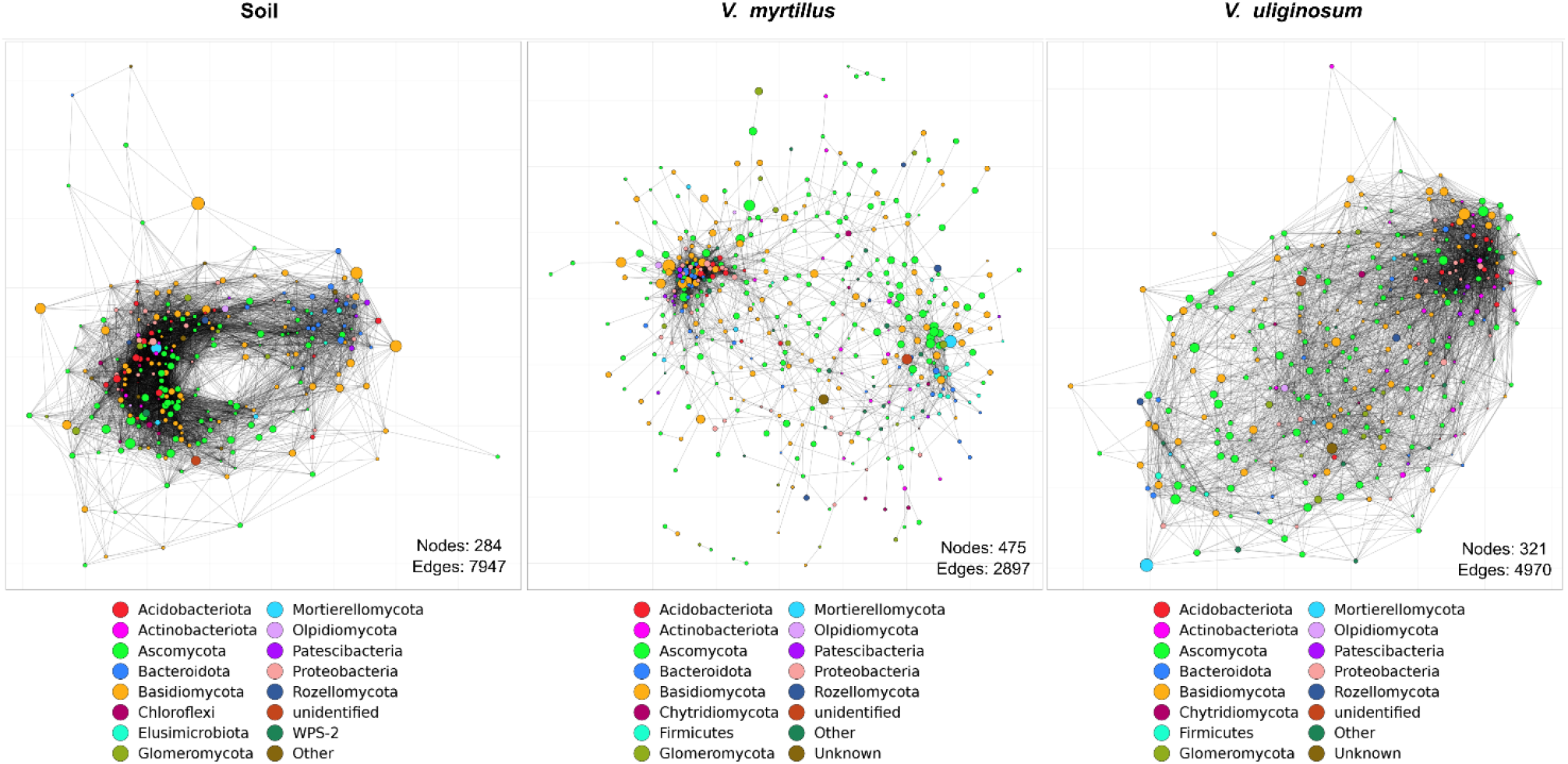
Co-occurrence networks strongly distinguish soil, *V myrtillus* and *V. uliginosum* microbiota. Network analysis of co-occurrence between bacterial and fungal phyla associated with *V. myrtillus* and *V. uliginosum* species and their soil. Each node in the network represents a phylum, and the connecting lines indicate positive co-occurrence interactions between ASVs affiliated to these phyla.

Concerning the two host plant species, the co-occurrence network of the microbial communities associated with *V. myrtillus* presents a higher number of nodes (475) than the soil or *V. uliginosum*, indicating a greater number of distinct microbial groups present within this plant species. However, the number of edges (2897) is lower than that of soil, suggesting that while *V. myrtillus* supports a diverse microbial community, the level of microbial interactions might be less complex than in the soil, possibly due to a more selective and specialized microbiome ^34,36^. The higher number of edges (4970), in *V. uliginosum* compared to *V. myrtillus,* together with the lower number of nodes (321), suggest that microbial taxa are more tightly ecologically connected in *V. uliginosum* than in *V. myrtillus*, with several taxa having the same co-occurrence patterns (lower number of nodes compared to *V. myrtillus*). This may imply that *V. uliginosum* microbiota could be more resilient to changes or that *V. uliginosum* better recruit soil taxa than *V. myrtillus*. However, we should be cautious to infer ecological properties of the microbiota from the sole network analysis ^37^.

### 3.4 Prediction of potential functions and metabolic pathways

Considering the taxonomic enrichment between host species and compartments, as well as differences in networks reported above, we hypothesized that some functional (metabolic and ecological) differences may be associated with such taxonomic variations. Consequently, functional profiles were inferred from retrieved taxonomies, in terms of the metabolic pathways and ecological roles present in the bacterial taxa (Figure 5) and trophic mode for fungi (Figure 6). Fermentation pathways showed higher relative abundance in *V. myrtillus* and *V. uliginosum*, especially in the root and leaf compartments. This suggests that endophytic microbiota of these plant species may rely on the use of unusual carbon sources, possibly including (e.g. in roots) anaerobic processes). Under this assumption, the aromatic compound degradation pathway, slightly more abundant in *V. uliginosum* leaves, is in line with the utilization of unusual carbon and energy sources by plant endophytes. Leaves of several plant species, especially medicinal plants, have shown the presence of a microbiota adapted to tolerate and thrive in presence of metabolites (often toxic) present in plant tissues ^38^. Since bilberry tissues harbor several compounds with antimicrobial activity ^39^ we cannot exclude that leaf microbiota of *V. myrtillus* and *V. uliginosum* could harbor metabolic activities able to tolerate and possibly utilize toxic compounds, including aromatic molecules, as carbon and energy sources. Another interesting function was the nitrate reduction pathway; this was notably more abundant in the roots of both *V. myrtillus* and *V. uliginosum*, than in other compartments, suggesting that plants, particularly *V. myrtillus*, host several bacteria which may help in soil nitrogen assimilation by roots. Actually, it is known that soil acidic pH reduces soil nitrate reduction potential ^40^, leading to lower nitrate levels. Additionally, acidic conditions favor ammonium (NH_4_^+^) retention over nitrate because nitrate is more prone to leaching in acidic soils ^41^. Acidophilic plants, as *V. myrtillus* and *V. uliginosum* may take advantage in recruiting beneficial nitrate reducing bacteria in their root apparatus. Among the other functions, the relative abundance of cellulolysis was highest in *V. uliginosum*, followed by *V. myrtillus*, and bulk soil. Notably, cellulolytic activity was also particularly pronounced in the rhizosphere and bulk soil. Cellulolytic activity could be related to the high abundance of Actinobacteria, which are known to harbor cellulase genes and are key players in organic matter cycling in the soil. We could then hypothesize that rhizosphere and soil have good potential for carbon cycling, then also humification processes. However, we cannot exclude that some cellulase activities, not detected in our analysis, could also be present in some plant pathogen. Actually, bacteria assigned as “animal parasites and symbionts” were found, especially prominent in *V. myrtillus* and *V. uliginosum*. Previous analyses have reported that other Vaccinium species also harbor potential symbiotic bacteria ^24^, as members of the order Rhizobiales, within the class of Alphaproteobacteria. However, we can likely assume that these bacteria behave as commensal endophytes, not as symbiotic nitrogen fixing bacteria (as they do with leguminous plants).

**Figure 5.**
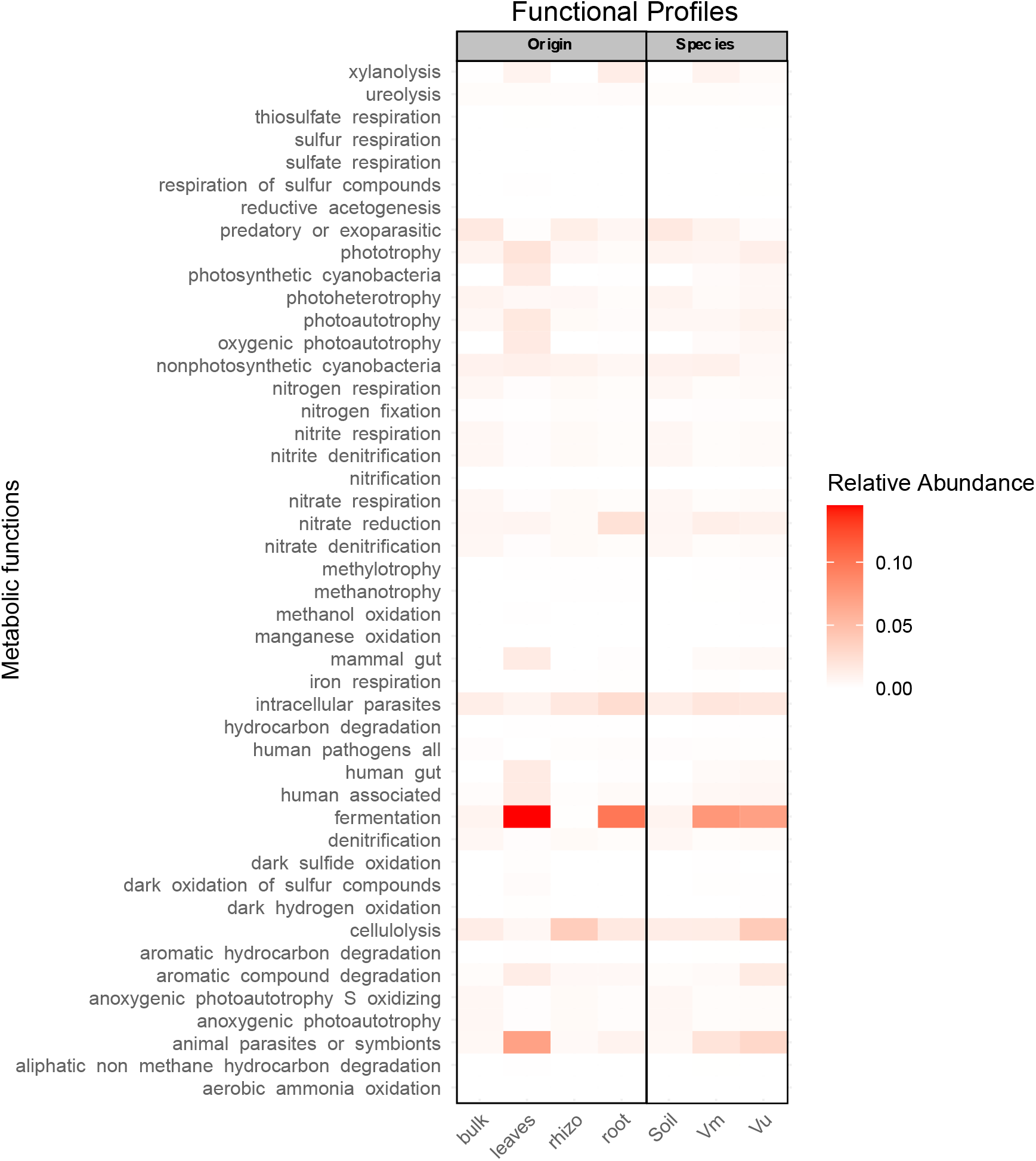
Metabolic pathways and their relative abundances across different conditions. The heatmap shows the relative abundances of ecologic functions and metabolic pathways identified through FAPROTAX in the bacterial microbiota. Abundances with respect to compartment (origin) and host the species are reported. (Vm, *V. myrtillus*; Vu, *V. uliginosum*; Soil, bulk soil. The color gradient indicates relative abundance, with darker shades representing higher values.

**Figure 6.**
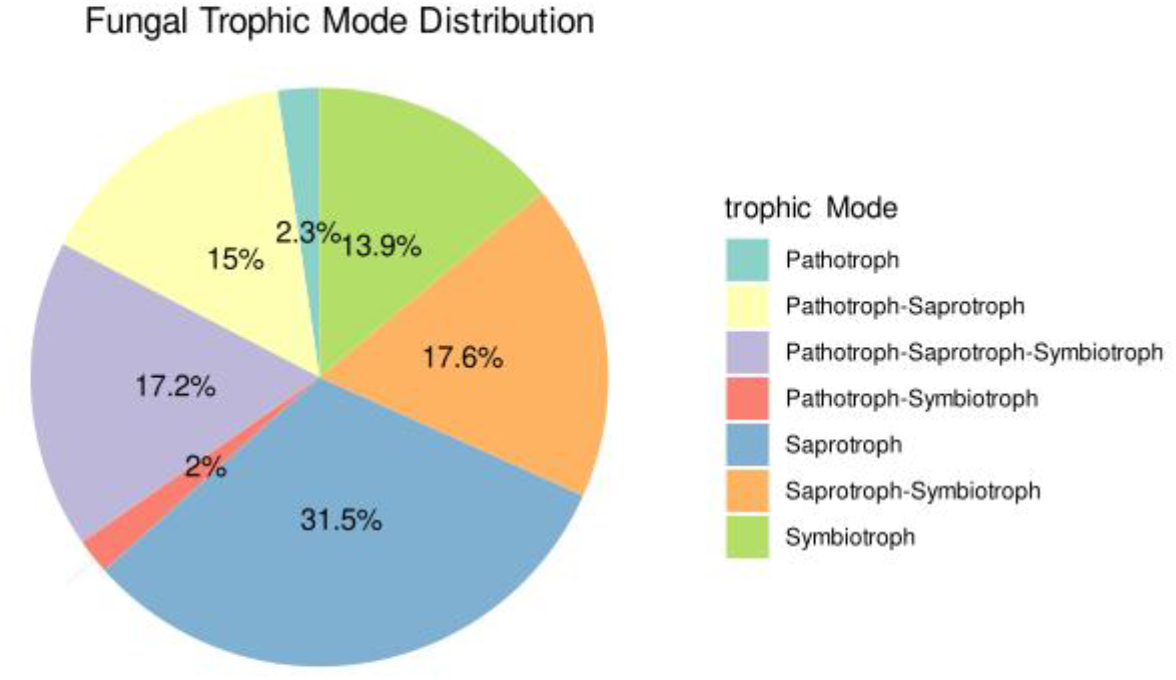
Pie chart of fungal trophic mode distribution, showing the relative abundance of each mode based on the total count. Only modes representing more than 0.5% are labeled.

The analysis of fungal trophic modes revealed a diverse distribution of functional roles within the community. The most abundant trophic mode is saprotroph, which accounts for 31.47% of the total occurrences, indicating that a significant portion of the fungal community is involved in decomposing dead organic matter, as previously observed for *V. myrtillus* in coniferous forests in Norther Europe ^32^. This is followed by saprotroph-symbiotroph (17.60%) and pathotroph-saprotroph-symbiotroph (17.16%), suggesting that many fungi exhibit mixed functional strategies, playing roles in both decomposition and symbiotic relationships with plants. Symbiotrophs (13.93%), which include ErM fungi ^42^, and pathotroph-saprotroph (14.97%) also contribute to the overall functional diversity, with fungi in these categories likely playing key roles in nutrient exchange and mutualistic symbioses. The relatively lower representation of pathotrophs (2.32%) and pathotroph-symbiotrophs (1.96%) indicates that pathogens, although present, are less prevalent in this community. This functional diversity highlights the complex ecological interactions within the fungal community, with a predominance of decomposers and symbionts contributing to nutrient cycling and plant interactions.

## 3. Conclusion

This study provides a comprehensive analysis of the bacterial and fungal microbiota associated with *V. myrtillus* and *V. uliginosum* in wild populations of the Central Italian Apennines. By utilizing 16S rRNA and ITS amplicon sequencing, we demonstrated that microbial communities differ significantly between plant compartments and species.

Our taxonomic analyses highlighted key microbial groups, including Alphaproteobacteria, Actinobacteria, and Acidobacteria among bacteria, and Agaricomycetes and Leotiomycetes among fungi. While these taxa have been found in other *Vaccinium* species, it is worth noticing that our study showed that *V. myrtillus* and *V. uliginosum* host different abundances of members of these classes. Our experimental plan cannot provide mechanistic indications on the role of such taxa in the biology of the two *Vaccinium* species, including the different colonization ability and possibly the production of the relevant secondary metabolites in leaves and fruits.

The co-occurrence network analysis supports the hypothesis that microbiota structure is shaped by both environmental and host-related factors. The interpretation on the number of nodes and edges identified suggests that the microbiota of *V. myrtillus* are more selectively filtered than that of *V. uliginosum*. We may speculate that *V. uliginosum* more tightly associated and possibly functionally integrated microbiota, makes the host plant potentially more resilient to environmental changes than *V. myrtillus*. Based on this hypothesis, it could be plausible that the invasiveness displayed by *V. uliginosum* populations against *V. myrtillus* populations could possibly relate to such a broader ability of the former species to recruit the soil microbiota.

The functional predictions, which indicated potential roles in nitrogen cycling, cellulose degradation, and plant-microbe interactions, with a particular emphasis on the microbial contributions to organic matter decomposition and nitrogen solubilization, may suggest several direct and indirect effects on plant species physiology and growth. However, the level of functional inference we may derive from 16S rRNA and ITS amplicon sequencing cannot provide evidence for differentiation between *V. myrtillus* and *V. uliginosum* microbiota at this level.

In conclusion, the findings obtained provide novel insights into the microbiota of wild bilberries, highlighting their ecological significance in subalpine environments. Understanding these microbial communities will be crucial for developing microbiome-based strategies to enhance plant resilience and productivity and develop innovative cultivation practices for *V. myrtillus*.

## 4. Materials and Methods

### 4.1 Sampling sites and sampling methods

Five sampling sites were identified in Central Apennines, Tuscany (Italy) (Table 2). In each site, populations of *V. myrtillus* were present. For two sites, coverage by *V. uliginosum* was also present, directly in contact with *V. myrtillus* vegetation. Sampling was performed on 29^th^ June 2023. Sampling methods were in accordance with standard methods defined for the Crop Microbiome Survey Initiative (https://www.globalsustainableagriculture.org/the-crop-microbiome-survey/) and previously published ^43^. Briefly, for each sampling site, a plot of 2 m x 2 m was identified, three randomly selected plants along the vertices of the plot were used to collect leaves, roots, and rhizospheric soil sample. Three bulk soil samples, at a depth of 5 cm, were also collected. The samples from individual plants and soil were mixed to produce a single composite sample. For each plot two duplicate composite samples were created.

**Table 2.**
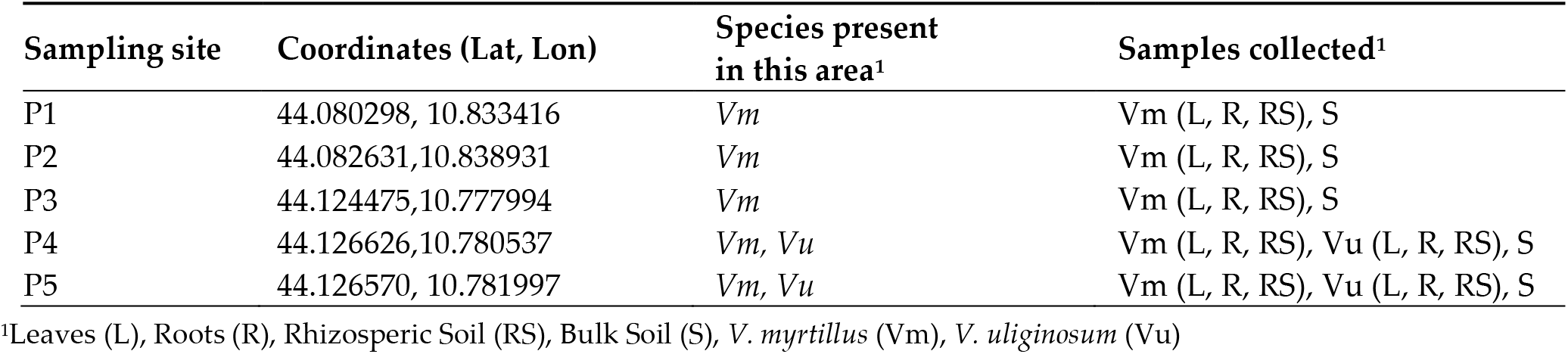
Description of the sampling sites. Acronyms are used to indicate Vaccinium species, plant compartment and soils collected.

| Sampling site | Coordinates (Lat, Lon) | Species present in this area<sup>1</sup> | Samples collected<sup>1</sup> |
| --- | --- | --- | --- |
| P1 | 44.080298, 10.833416 | *Vm* | *Vm* (L, R, RS), S |
| P2 | 44.082631, 10.838931 | *Vm* | *Vm* (L, R, RS), S |
| P3 | 44.124475, 10.777994 | *Vm* | *Vm* (L, R, RS), S |
| P4 | 44.126626, 10.780537 | *Vm*, *Vu* | *Vm* (L, R, RS), *Vu* (L, R, RS), S |
| P5 | 44.126570, 10.781997 | *Vm*, *Vu* | *Vm* (L, R, RS), *Vu* (L, R, RS), S |

<sup>1</sup>Leaves (L), Roots (R), Rhizospheric Soil (RS), Bulk Soil (S), *V. myrtillus* (Vm), *V. uliginosum* (Vu)

### 4.2 eDNA extraction and amplicon sequencing

Environmental DNA (eDNA) was extracted from 250 mg of soils and plant tissues using the DNeasyPowerSoil Pro (Qiagen Italy, Milan, Italy). The V3-V4 region of the bacterial 16SrRNA gene was amplified as reported in the Illumina 16S Metagenomic Sequencing Library Preparation protocol with primers as reported in ^44^. Fungal Internal Transcribed Spacer 1 (ITS1) was amplified following standards of Hearth Microbiome Project (https://earthmicrobiome.org/protocols-and-standards/its/) and ITS1f-ITS2 primer pairs ^45^. Amplicons were used to construct libraries and sequenced on an Illumina NovaSeq6000 instrument (Illumina, San Diego, CA) SP flow cell using 250 bp paired-end cycles at Biomarker Technologies (BMK) GmbH (Münster, Germany), following Illumina protocols. Reads were trimmed using Cutadapt ^46^ to remove the following sequencing primers: V3-V4_F: ACTCCTACGGGAGGCAGCA, V3-V4_R: GGACTACHVGGGTWTCTAAT, ITS1_F: CTTGGTCATTTAGAGGAAGTAA, ITS1_R: GCTGCGTTCTTCATCGATGC. Raw reads are available from NCBI SRA database under Bioproject PRJNA1250273.

### 4.3 Bioinformatic analysis of sequencing data, clustering of reads and taxonomic assignment

Data analysis was performed as previously reported ^47^. In brief, DADA2 pipeline (version 1.24.0) ^48^ as used to cluster amplicon sequence variants (ASVs). Bacterial taxonomy assignment was carried out comparing 16SrRNA ASV against SILVA_SSU_r138 database ^49^ using “DECIPHER” R package (version 2.24.1) ^50^. Annotated ASVs count tables were processed in Phyloseq package ^51^. All sequences classified as chloroplasts were removed. For ITS ASV taxonomy assignment was carried out on the UNITE ITS database ^52^, (UNITE_v2020_February2020).

### 4.4 Statistical analysis of microbiota diversity

Alpha diversity (Shannon and Pielou’s Evenness indices) were calculated using the function “diversity()” within “Phyloseq” R package ^51^. Good’s coverage and evenness indices were calculated through the R functions ‘goods()’ and ‘evenness()’, respectively, within the ‘microbiome’ R package (version 1.12.0). Taxonomic differences among microbiota were inspected by non-metric multidimensional scaling (nMDS), using the “ordinate” function and plotting by the “plot_ordination()” function within phyloseq package. The “ggplot2” R package (version 3.3.6) ^53^ was used to generate relative abundance plots. Different community structures were analyzed using permutational a multivariate analysis of variance (PERMANOVA) using the R packages “vegan” (version 2.6.2) and the function “adonis2()”. Wilcoxon tests for multiple comparisons of averages were performed on alpha diversity indices using ‘ggviolin()’ and ‘stat_compare_means()’ within the ‘ggpubr’ R package (version 0.4.0).

Differential abundance analysis was performed using the R package DESeq2 (version 1.36.0) ^54^. Network analysis of ASVs co-occurrence was performed using SpeSpeNet ^55^.

### 4.5 Functional inference on bacterial and fungal microbiota

Ecological functions of microbial communities were predicted with FAPROTAX ^56^ for 16S sequencing data, while for the fungal microbiome, the ecological roles were assigned using”FunguildR” package (version 0.2.0.900) ^57^.

## Supplementary Materials

The following supporting information can be downloaded at: www.mdpi.com/xxx/s1 Supplementary Information File; Supplementary Dataset.

## Author Contributions

Conceptualization, E.G., A.M., F.V., and C.F.; methodology, F.V.; validation, F.V., C.F., E.G. S.B., and A.M.; formal analysis, G.C., F.V., and C.F.; investigation, F.V., C.F., G.C.; resources, A.M. E.G. and S.B.; data curation, F.V., C.F.; writing—original draft preparation, F.V., A.M. and C.F.; writing—review and editing, F.V. A.M. C.F. E.G., and S.B.; visualization, F.V.; supervision, A.M.; project administration, A.M., C.F.; funding acquisition, A.M.. All authors have read and agreed to the published version of the manuscript.

## Data Availability Statement

Sequence reads are available on SRA database under the Bioproject PRJNA1250273.

## Abbreviations

The following abbreviations are used in this manuscript:

16SrRNA: 16S ribosomal RNA

ITS: Internal Transcribed spacer

nMDS: nonmetric Multimensional scaling

