## Supplementary material for "Comparative Analysis of the Microbiota of *Vaccinium myrtillus* and *Vaccinium uliginosum* in the Central Italian Apennines": Supplementary Information File Vaccaro et al.pdf

**Table S1:** V3-V4 region of 16S rRNA reads passing quality filtering and metadata. In order: input (raw sequences obtained directly from the sequencing machine), filtered (low-quality reads are removed), denoisedF and denoisedR (forward and reverse sequences after error correction), merged and nonchim (chimeric sequences are removed). Percentages reported in the last row refer to the initial number of input read.

| | **input** | **filtered** | **denoisedF** | **denoisedR** | **merged** | **nonchim** |
|---|---|---|---|---|---|---|
| **M1-1** | 79804 | 72994 | 69986 | 69664 | 36275 | 33456 |
| **M1-2** | 79980 | 74082 | 70709 | 70190 | 34060 | 31065 |
| **M10-1** | 80000 | 73459 | 70151 | 70112 | 39178 | 37273 |
| **M10-2** | 80024 | 74052 | 71116 | 70739 | 40562 | 38764 |
| **M11-1** | 79707 | 73511 | 71429 | 71565 | 42895 | 41457 |
| **M11-2** | 79752 | 73486 | 71117 | 71202 | 42457 | 40930 |
| **M12-1** | 79826 | 73605 | 73542 | 73399 | 63664 | 63487 |
| **M12-2** | 80038 | 74183 | 74124 | 74099 | 64562 | 64462 |
| **M13-1** | 80030 | 73347 | 70719 | 70760 | 45083 | 43278 |
| **M13-2** | 80109 | 74478 | 72489 | 72761 | 48032 | 46114 |
| **M14-1** | 79728 | 73979 | 72313 | 72443 | 46576 | 45036 |
| **M14-2** | 79928 | 73747 | 72483 | 72610 | 45599 | 44162 |
| **M15-1** | 79889 | 73803 | 73205 | 73290 | 64453 | 63434 |
| **M15-2** | 80140 | 73343 | 73041 | 73188 | 64700 | 63377 |
| **M16-1** | 79976 | 73590 | 72032 | 71901 | 57858 | 18490 |
| **M16-2** | 79955 | 73251 | 71349 | 71604 | 57527 | 23361 |
| **M17-1** | 79740 | 73877 | 70888 | 70526 | 41828 | 39851 |
| **M17-2** | 79932 | 72994 | 70319 | 69947 | 41439 | 39450 |
| **M18-1** | 80117 | 73914 | 71757 | 71561 | 51745 | 50140 |
| **M18-2** | 80155 | 73456 | 72801 | 72930 | 58882 | 49032 |
| **M19-1** | 79835 | 73639 | 72918 | 73031 | 51537 | 37575 |
| **M19-2** | 80065 | 73430 | 73253 | 73225 | 52283 | 46169 |
| **M2-1** | 79932 | 73662 | 70645 | 70434 | 38152 | 35987 |
| **M2-2** | 80227 | 74697 | 71813 | 71500 | 38717 | 36654 |
| **M20-1** | 79981 | 74319 | 71921 | 71750 | 43545 | 41696 |
| **M20-2** | 80242 | 73699 | 71360 | 71040 | 42220 | 40630 |
| **M21-1** | 80181 | 73788 | 71618 | 71248 | 48271 | 46702 |

| | | | | | | |
|---|---|---|---|---|---|---|
| **M21-2** | 79884 | 73713 | 71345 | 71418 | 47004 | 45453 |
| **M22-1** | 80329 | 74044 | 72500 | 72621 | 50583 | 49399 |
| **M22-2** | 80172 | 74097 | 72897 | 72844 | 50424 | 49219 |
| **M23-1** | 79996 | 73845 | 73488 | 73454 | 47512 | 41895 |
| **M23-2** | 79960 | 73475 | 71512 | 71909 | 50510 | 19999 |
| **M24-1** | 79878 | 74009 | 73165 | 73237 | 51263 | 50030 |
| **M24-2** | 79920 | 74016 | 73045 | 73121 | 51742 | 50685 |
| **M25-1** | 80069 | 74092 | 71584 | 71715 | 50533 | 48670 |
| **M25-2** | 80151 | 73692 | 71744 | 72227 | 52090 | 31981 |
| **M26-1** | 80091 | 73712 | 71360 | 71873 | 30815 | 26218 |
| **M26-2** | 80146 | 72959 | 72691 | 72680 | 43142 | 37529 |
| **M3-1** | 80027 | 74190 | 72067 | 72015 | 46018 | 44586 |
| **M3-2** | 80262 | 73767 | 72120 | 72514 | 46729 | 45349 |
| **M4-1** | 79922 | 73676 | 72378 | 72203 | 55369 | 30200 |
| **M4-2** | 79930 | 73548 | 73066 | 73147 | 57980 | 53528 |
| **M5-1** | 79972 | 74218 | 70010 | 70414 | 36985 | 31228 |
| **M5-2** | 80047 | 73600 | 71503 | 71560 | 40221 | 38336 |
| **M6-1** | 79982 | 73676 | 71978 | 72057 | 40206 | 38646 |
| **M6-2** | 80237 | 73728 | 71880 | 71824 | 41419 | 39922 |
| **M7-1** | 80274 | 73843 | 72528 | 72588 | 44270 | 42635 |
| **M7-2** | 79608 | 73744 | 71794 | 71652 | 43039 | 41491 |
| **M8-1** | 79818 | 73564 | 73109 | 73088 | 49200 | 42866 |
| **M8-2** | 79918 | 74461 | 74308 | 74249 | 54002 | 48813 |
| **M9-1** | 79981 | 74221 | 71289 | 71173 | 43816 | 41865 |
| **M9-2** | 80100 | 74266 | 71213 | 71228 | 43904 | 41798 |
| **Sum** | 4159967 | 3836541 | 3743672 | 3743530 | 2470876 | 2194373 |
| % | | 92,2 | 89,9 | 89,9 | 59,4 | 52,7 |

**Table S2:** ITS reads passing quality filtering and metadata. In order: input (raw sequences obtained directly from the sequencing machine), filtered (low-quality reads are removed), denoisedF and denoisedR (forward and reverse sequences after error correction), merged and nonchim (chimeric sequences are removed). Percentages reported in the last row refer to the initial number of input read.

| | **input** | **filtered** | **denoisedF** | **denoisedR** | **merged** | **nonchim** |
|---|---|---|---|---|---|---|
| **M1-1** | 80241 | 68464 | 67130 | 66883 | 64757 | 64647 |
| **M1-2** | 79895 | 66367 | 65899 | 65971 | 65182 | 65180 |
| **M10-1** | 79797 | 73443 | 73008 | 73036 | 70811 | 70811 |
| **M10-2** | 80178 | 73363 | 73094 | 73119 | 71914 | 71886 |
| **M11-1** | 80029 | 72377 | 72171 | 72221 | 69211 | 69192 |
| **M11-2** | 79853 | 72593 | 72309 | 72262 | 70492 | 70466 |
| **M12-1** | 80063 | 72107 | 70827 | 70991 | 64882 | 64027 |
| **M12-2** | 80388 | 73701 | 73610 | 73653 | 71387 | 70978 |
| **M13-1** | 80040 | 70971 | 70624 | 70728 | 67688 | 67446 |
| **M13-2** | 80330 | 70839 | 70335 | 70530 | 68749 | 68496 |
| **M14-1** | 79986 | 72729 | 72585 | 72574 | 71631 | 71193 |
| **M14-2** | 80136 | 71656 | 71500 | 71470 | 67897 | 67705 |
| **M15-1** | 80060 | 72271 | 72237 | 72211 | 71769 | 71769 |
| **M15-2** | 80010 | 71622 | 71604 | 71602 | 70933 | 70200 |
| **M16-1** | 79825 | 71552 | 71435 | 71478 | 70779 | 69884 |
| **M16-2** | 79995 | 72158 | 72114 | 72091 | 70998 | 70601 |
| **M17-1** | 80016 | 71171 | 70529 | 70590 | 61589 | 61366 |

| | | | | | | |
|---|---|---|---|---|---|---|
| **M17-2** | 80124 | 71357 | 70954 | 70838 | 62878 | 62766 |
| **M18-1** | 80348 | 72741 | 72142 | 72163 | 65257 | 64284 |
| **M18-2** | 80009 | 72179 | 72012 | 71997 | 70846 | 70842 |
| **M19-1** | 79929 | 73003 | 72869 | 72920 | 72650 | 72308 |
| **M19-2** | 79959 | 71813 | 71674 | 71699 | 71247 | 70746 |
| **M2-1** | 80165 | 72823 | 72435 | 72428 | 70962 | 70236 |
| **M2-2** | 79938 | 72425 | 72107 | 72166 | 70446 | 66530 |
| **M20-1** | 79884 | 72626 | 72428 | 72461 | 71467 | 71344 |
| **M20-2** | 80223 | 72771 | 72431 | 72468 | 65560 | 65549 |
| **M21-1** | 80010 | 71632 | 70926 | 70978 | 65926 | 65228 |
| **M21-2** | 79866 | 73144 | 72865 | 72855 | 70416 | 70060 |
| **M22-1** | 79938 | 72470 | 72401 | 72362 | 67522 | 67210 |
| **M22-2** | 79996 | 73003 | 72764 | 72848 | 69199 | 69011 |
| **M23-1** | 56963 | 50402 | 50303 | 50366 | 50007 | 50007 |
| **M23-2** | 80000 | 71078 | 70881 | 71048 | 70607 | 70500 |
| **M24-1** | 79911 | 71918 | 71037 | 71269 | 65061 | 64093 |
| **M24-2** | 80249 | 73216 | 73065 | 73058 | 68862 | 68794 |
| **M25-1** | 80084 | 72516 | 72221 | 72376 | 64409 | 64254 |
| **M25-2** | 79870 | 70942 | 69903 | 70043 | 64992 | 64215 |
| **M26-1** | 80091 | 71474 | 71310 | 71444 | 69912 | 69498 |
| **M26-2** | 80039 | 72295 | 71920 | 71774 | 68023 | 66956 |
| **M3-1** | 80199 | 72179 | 71799 | 71944 | 69224 | 69147 |
| **M3-2** | 79870 | 71827 | 71445 | 71584 | 69007 | 68918 |
| **M4-1** | 80031 | 69692 | 68934 | 68804 | 66501 | 66267 |
| **M4-2** | 79744 | 72338 | 72263 | 72194 | 71800 | 70531 |
| **M5-1** | 79933 | 72907 | 72660 | 72675 | 71188 | 70774 |
| **M5-2** | 79945 | 73130 | 72923 | 72964 | 72031 | 70608 |
| **M6-1** | 80174 | 73341 | 72969 | 73041 | 70113 | 69723 |
| **M6-2** | 80080 | 72864 | 72669 | 72703 | 69830 | 69243 |
| **M7-1** | 80335 | 72591 | 72540 | 72522 | 69584 | 69255 |
| **M7-2** | 79983 | 72987 | 72893 | 72888 | 71278 | 71097 |
| **M8-1** | 79799 | 73117 | 73087 | 73092 | 72818 | 72179 |
| **M8-2** | 79935 | 73904 | 73841 | 73812 | 72116 | 71178 |
| **M9-1** | 79901 | 71530 | 70105 | 70193 | 64488 | 64121 |
| **M9-2** | 79916 | 71640 | 71029 | 71131 | 65325 | 64521 |
| **Sum** | 4138283 | 3727259 | 3708816 | 3710518 | 3562221 | 3537840 |
| % | | 90,1 | 89,6 | 89,7 | 86,1 | 85,5 |

a)

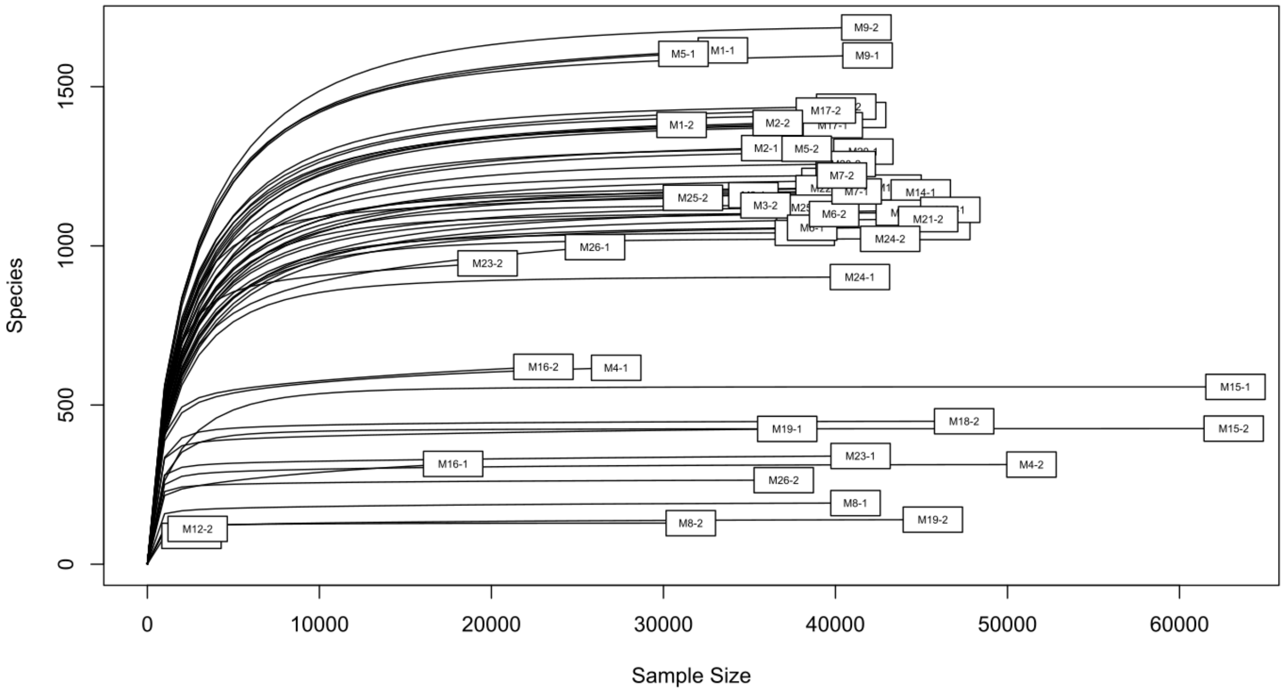

b)

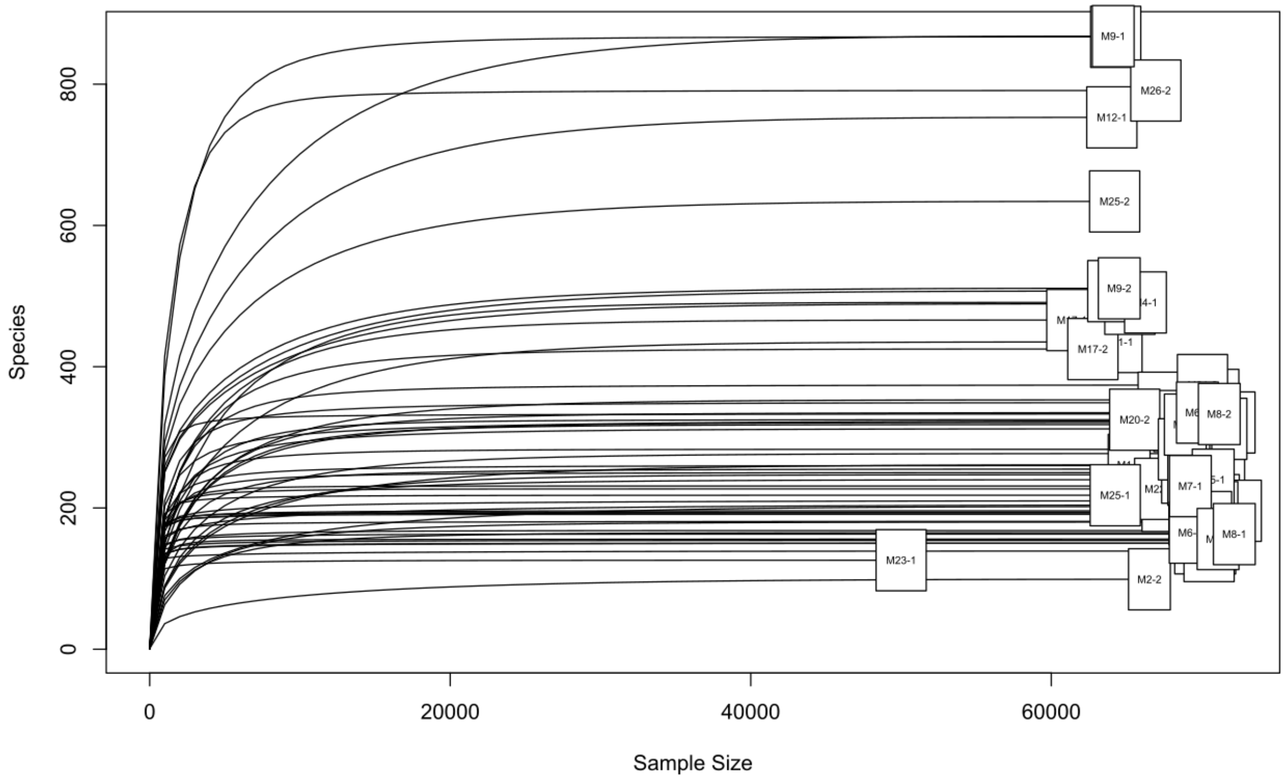

**Figure S1:** Rarefaction curve of all samples, the y axis indicated the species richness while the x axis indicates the sequence sample size, the curves reached a plateau for all samples, indicating a satisfactory survey of the bacterial diversity. Rarefaction curves were obtained from V3-V4 amplicon sequencing (a) and ITS sequencing (b).

**Table S3:** Goods Coverage for V3-V4 sequencing.

| | no.sing | no.seqs | goods |
|---|---|---|---|
| **M1-1** | 83 | 33453 | 99.7518907123 |
| **M1-2** | 62 | 31056 | 99.8003606388 |
| **M10-1** | 31 | 37265 | 99.916812022 |
| **M10-2** | 42 | 38757 | 99.8916324793 |
| **M11-1** | 29 | 41201 | 99.9296133589 |
| **M11-2** | 28 | 40624 | 99.9310752265 |
| **M12-1** | 0 | 2565 | 100 |
| **M12-2** | 2 | 2923 | 99.9315771468 |
| **M13-1** | 30 | 43262 | 99.9306550784 |
| **M13-2** | 20 | 46095 | 99.9566113461 |
| **M14-1** | 17 | 44959 | 99.9621877711 |
| **M14-2** | 10 | 44086 | 99.9773170621 |
| **M15-1** | 0 | 63252 | 100 |
| **M15-2** | 1 | 63150 | 99.9984164687 |
| **M16-1** | 47 | 17776 | 99.7355985599 |
| **M16-2** | 57 | 23025 | 99.7524429967 |
| **M17-1** | 31 | 39844 | 99.9221965666 |
| **M17-2** | 38 | 39439 | 99.9036486726 |
| **M18-1** | 27 | 39764 | 99.9320993864 |
| **M18-2** | 2 | 47480 | 99.9957877001 |
| **M19-1** | 20 | 37196 | 99.9462307775 |
| **M19-2** | 4 | 45660 | 99.991239597 |
| **M2-1** | 45 | 35978 | 99.8749235644 |
| **M2-2** | 51 | 36646 | 99.86083065 |
| **M20-1** | 31 | 41621 | 99.9255183681 |
| **M20-2** | 24 | 40595 | 99.9408794186 |
| **M21-1** | 35 | 46672 | 99.9250085704 |
| **M21-2** | 16 | 45388 | 99.9647483916 |
| **M22-1** | 17 | 39423 | 99.9568779646 |
| **M22-2** | 9 | 38219 | 99.9764515032 |
| **M23-1** | 12 | 41460 | 99.9710564399 |
| **M23-2** | 63 | 19777 | 99.6814481468 |
| **M24-1** | 6 | 41421 | 99.985514594 |
| **M24-2** | 6 | 43180 | 99.9861046781 |
| **M25-1** | 21 | 38303 | 99.9451740073 |
| **M25-2** | 38 | 31713 | 99.8801753224 |
| **M26-1** | 108 | 26022 | 99.5849665668 |
| **M26-2** | 4 | 36993 | 99.9891871435 |
| **M3-1** | 33 | 35230 | 99.9063298325 |
| **M3-2** | 12 | 35936 | 99.9666073019 |
| **M4-1** | 41 | 27252 | 99.8495523264 |
| **M4-2** | 2 | 51395 | 99.9961085709 |
| **M5-1** | 81 | 31165 | 99.7400930531 |
| **M5-2** | 29 | 38327 | 99.924335325 |
| **M6-1** | 20 | 38646 | 99.9482482016 |
| **M6-2** | 16 | 39922 | 99.9599218476 |
| **M7-1** | 15 | 41231 | 99.9636196066 |
| **M7-2** | 18 | 40374 | 99.9554168524 |
| **M8-1** | 4 | 41164 | 99.9902827714 |
| **M8-2** | 0 | 31590 | 100 |
| **M9-1** | 39 | 41865 | 99.9068434253 |

| | | | |
|---|---|---|---|
| **M9-2** | 30 | 41798 | 99.9282262309 |
| **Mean** | | | 99.9161893124 |

**Table S4:** Goods Coverage for ITS sequencing.

| | no.sing | no.seqs | goods |
|---|---|---|---|
| **M1-1** | 0 | 64647 | 100 |
| **M1-2** | 0 | 65180 | 100 |
| **M10-1** | 0 | 70811 | 100 |
| **M10-2** | 0 | 71886 | 100 |
| **M11-1** | 0 | 69192 | 100 |
| **M11-2** | 0 | 70466 | 100 |
| **M12-1** | 0 | 64027 | 100 |
| **M12-2** | 0 | 70978 | 100 |
| **M13-1** | 0 | 67446 | 100 |
| **M13-2** | 0 | 68496 | 100 |
| **M14-1** | 0 | 71193 | 100 |
| **M14-2** | 0 | 67705 | 100 |
| **M15-1** | 0 | 71769 | 100 |
| **M15-2** | 0 | 70200 | 100 |
| **M16-1** | 0 | 69884 | 100 |
| **M16-2** | 0 | 70601 | 100 |
| **M17-1** | 0 | 61366 | 100 |
| **M17-2** | 0 | 62766 | 100 |
| **M18-1** | 0 | 64284 | 100 |
| **M18-2** | 0 | 70842 | 100 |
| **M19-1** | 0 | 72308 | 100 |
| **M19-2** | 0 | 70746 | 100 |
| **M2-1** | 0 | 70236 | 100 |
| **M2-2** | 0 | 66530 | 100 |
| **M20-1** | 0 | 71344 | 100 |
| **M20-2** | 0 | 65549 | 100 |
| **M21-1** | 0 | 65228 | 100 |
| **M21-2** | 1 | 70060 | 99.998572652 |
| **M22-1** | 0 | 67210 | 100 |
| **M22-2** | 0 | 69011 | 100 |
| **M23-1** | 0 | 50007 | 100 |
| **M23-2** | 0 | 70500 | 100 |
| **M24-1** | 0 | 64093 | 100 |
| **M24-2** | 0 | 68794 | 100 |
| **M25-1** | 0 | 64254 | 100 |
| **M25-2** | 0 | 64215 | 100 |
| **M26-1** | 0 | 69498 | 100 |
| **M26-2** | 0 | 66956 | 100 |
| **M3-1** | 0 | 69147 | 100 |
| **M3-2** | 0 | 68918 | 100 |
| **M4-1** | 0 | 66267 | 100 |
| **M4-2** | 0 | 70531 | 100 |
| **M5-1** | 0 | 70774 | 100 |
| **M5-2** | 0 | 70608 | 100 |
| **M6-1** | 1 | 69723 | 99.9985657531 |
| **M6-2** | 0 | 69243 | 100 |
| **M7-1** | 0 | 69255 | 100 |
| **M7-2** | 0 | 71097 | 100 |
| **M8-1** | 0 | 72179 | 100 |
| **M8-2** | 0 | 71178 | 100 |
| **M9-1** | 0 | 64121 | 100 |

| | | | |
|---|---|---|---|
| **M9-2** | 0 | 64521 | 100 |
| **Mean** | | | 99.9999449693 |
